# Implicit semantic prediction error can account for N400 effects on articles that do not differ in meaning: A neural network model

**DOI:** 10.1101/569020

**Authors:** Milena Rabovsky

## Abstract

N400 effects on indefinite articles (a/an) compatible or incompatible with expected nouns have been initially taken as strong evidence for probabilistic pre-activation of phonological word forms, and recently been intensely debated because they have been difficult to replicate. Here, we simulate these effects using a neural network model of sentence comprehension that we previously used to simulate a broad range of empirical N400 effects. The model produces the effects when the cue validity of the articles concerning upcoming noun meaning is high, but fails to produce the effects when the cue validity of the articles is low due to adjectives presented between articles and nouns during training, providing a possible explanation for the small size of the effects in empirical studies. The model accounts for article induced N400 effects without assuming pre-activation of word forms, and instead simulates these effects as the stimulus-induced change in a probabilistic representation of meaning corresponding to an implicit semantic prediction error.

## Introduction

The N400 component of the event-related brain potential (ERP) has received a great deal of attention because it provides an electrophysiological indicator of meaning processing in the brain (Kutas & Federmeier, 2011). One issue that has been addressed using the N400 component and has triggered intense debates is the question in how far upcoming input is predicted and thus pre-activated during language comprehension. Specifically, in a landmark study DeLong and colleagues obtained larger N400 amplitudes to articles incompatible with an expected noun such as e.g., ‘The day was breezy so the boy went outside to fly an…’ where ‘an’ is incompatible with the expected continuation ‘kite’ (DeLong, Urbach, & Kutas, 2005). This result was taken to provide clear evidence for predictive pre-activation in language comprehension. Furthermore, because ‘a’ and ‘an’ arguably do not differ in meaning, it was taken to suggest predictive pre-activation of the form of upcoming words.

Earlier studies had already shown that N400 amplitudes are reliably reduced for predictable language input (see Kutas & Federmeier, 2011, for a review) such as for instance demonstrated by influences of cloze probability, which refers to the percentage of participants continuing a sentence fragment with a specific word in offline sentence completion tasks. For example, N400 amplitudes are smaller for high cloze probability continuations such as “Don’t touch the wet *paint*” as compared to low cloze probability continuations such as “Don’t touch the wet *dog*” (Kutas & Hillyard, 1984). However, in most studies, it is difficult to unequivocally decide whether reduced N400 amplitudes reflect facilitated processing due to prediction/ pre-activation of upcoming input (Kutas & Federmeier, 2000), or whether reduced N400 amplitudes reflect facilitated bottom-up processing because the incoming input better fits and is thus easier to integrate into the preceding context (Brown & Hagoort, 1993). Traditionally, the pre-activation account has been more closely linked to word-level processing (pre-activation of word representations) and the bottom up account has been more closely linked to sentence level processing (integration of a word into the sentence context).

However, pre-activation does not need to be restricted to word level representations as demonstrated for instance by our recent study simulating a broad range of 16 empirically observed N400 effects by treating N400 amplitudes as the change in a neural network model’s hidden layer activation state representing predicted sentence meaning (Rabovsky, Hansen, & McClelland, 2018; see also Baggio & Hagoort, 2011; Kutas & Federmeier, 2011, for other accounts going beyond the strict separation of word and sentence processing in explaining the N400). In this view, N400 amplitudes reflect an implicit prediction error or Bayesian surprise at the level of integrated (sentence) meaning, which is assumed to be predicted and thus pre-activated based on the experience of statistical regularities in the environment (see also Kuperberg & Jaeger, 2016; Rabovsky & McRae, 2014). Importantly, N400 effects induced by articles that do not differ in meaning but just in their form (i.e., ‘a’ and ‘an’) cannot be explained by differences in bottom-up integration difficulty, as they should be equally easy to integrate into the semantic context. Therefore, the observed N400 effects on articles have been taken as strong evidence for pre-activation of upcoming input during language comprehension (DeLong et al., 2005).

Pre-activation of which aspects of the upcoming input is reflected in these article-induced N400 effects? As noted above, DeLong and colleagues originally took the effects to reflect prediction of the phonological form of the next word, based on the reasoning that this phonological form, i.e., whether the next noun starts with a consonant or a vowel, determines whether the indefinite article should be ‘a’ or ‘an’ so that encountering the articles allows to evaluate whether the phonological form of the next word was predicted correctly. Following this and similar reasoning, article induced N400 effects have often been cited as evidence for prediction of upcoming language input at the level of word forms (e.g., Altmann & Mirković, 2009; Hagoort, 2017; Lau, Phillips, & Poeppel, 2008; Pickering & Garrod, 2013). However, an alternative perspective, which corresponds to the perspective implemented in our model, is that the article-induced N400 effects do not reflect prediction of word forms, bur rather reflect the change in a probabilistic representation of sentence meaning, which is cued by encountering the articles (see Fig. 1). From this view, predictions are not explicitly evaluated, but rather adjusted based on new incoming evidence, and N400 amplitudes reflect the shift in the conditional probabilities of semantic features cued by this new incoming evidence, i.e., the articles. The main goal of the current study is to demonstrate via explicit simulations how this perspective can mechanistically account for article induced N400 effects, thus offering an alternative to the original and still very common interpretation of the empirical effects.

**Figure 1.**
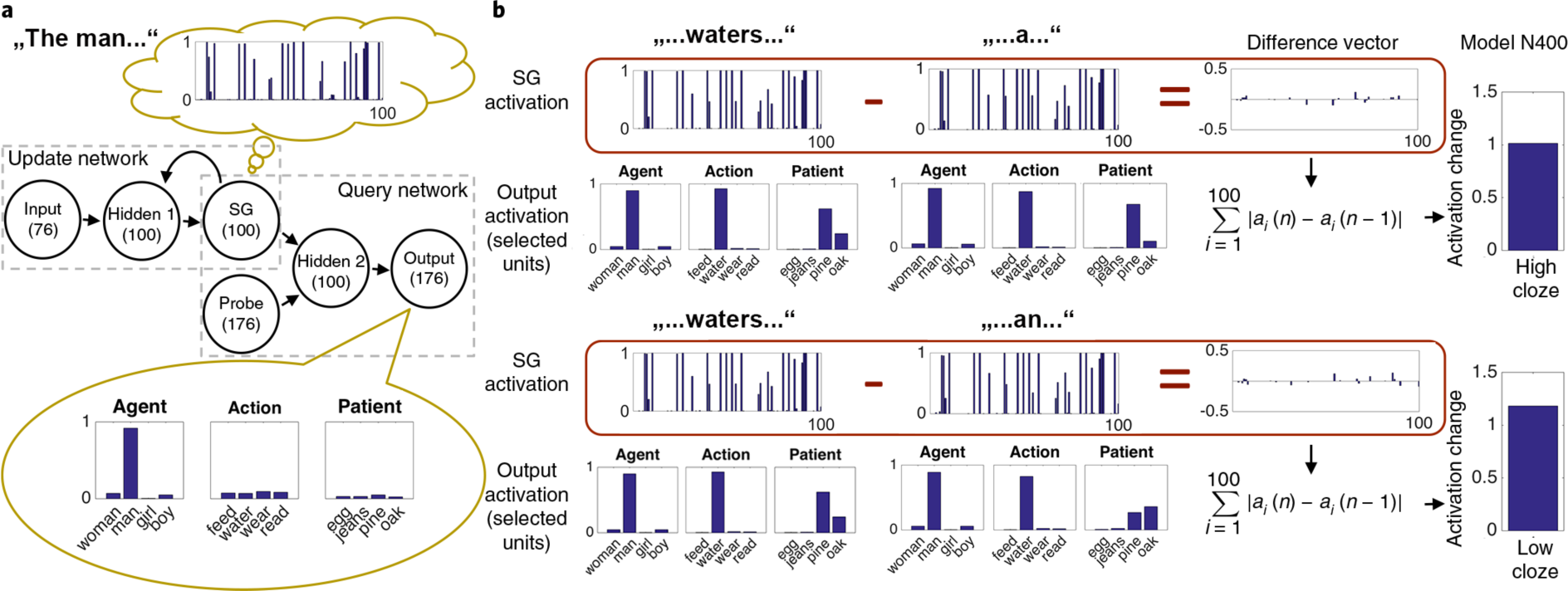
The Sentence Gestalt (SG) model architecture, processing a sentence with a high or low cloze probability article, and the model’s N400 correlate. The model (left) consists of an update network and a query network. Ovals represent layers of units (and number of units in each layer). Arrows represent all-to-all modifiable connections; each unit applies a sigmoid transformation to its summed inputs, where each input is the product of the activation of the sending unit times the weight of that connection. In the update part of the model, each incoming word is processed through layer Hidden 1 where it combines with the previous SG activation to produce the updated SG activation corresponding to the updated implicit representation of the described event. During training, after each presented word, the model is probed and given feedback concerning all aspects of the described event (e.g. agent, “man”, action, “play”, etc.) in the query network. Here, the activation from the probe layer combines via layer Hidden 2 with the current SG pattern to produce output activations. Selected output units activated in response to the agent, action, and patient probes are shown; each query response includes a distinguishing feature (e.g. ‘man’, ‘woman’, as shown) as well as other features (e.g., ‘person’, ‘adult’, not shown) that capture semantic similarities among event participants). After presentation of “The man”, the SG representation (thought bubble at top left) supports activation of the correct features when probed for the agent and estimates the probabilities of action and patient features. After the word „waters” (shown twice in the middle) the SG representation is updated and the model now activates the correct features given the agent and action probes, and estimates the probability of alternative possible patients, based on its experience (the man waters pines more often than oaks). If the next word is „a” (top), which is compatible with the high cloze probability continuation „pine“, the change in SG activation (summed magnitudes of changes in ‘Difference vector’) is smaller than if the next word is „an” (bottom), which is incompatible with „pine” and instead to be followed by the low cloze probability continuation „oak”. The change, called Semantic Update (SU) is the proposed N400 correlate (right). It is larger for the article compatible with the less as compared to the more probable described event.

Another issue concerning article induced N400 effects, which has recently been intensely debated, is that these effects have not always been replicated (for discussion see DeLong, Urbach, & Kutas, 2017; Ito, Martin, & Nieuwland, 2017b, 2017a), and in any case seem to be considerably smaller than observed in the original study (Nicenboim, Vasishth, & Rösler, 2019; Nieuwland et al., 2018). It has been suggested that this might be due to the fact that articles do not deterministically predict specific nouns in natural language (Ito et al., 2017b; Nieuwland et al., 2018). A second goal of the current study was to illustrate the importance of this factor, i.e. the cue validity of the articles concerning upcoming meaning, by including one simulation where articles deterministically predict the upcoming nouns during training (Simulation 1) and one simulation where this predictive relationship vanishes due to adjectives presented between articles and nouns during training (in analogy to, e.g., ‘an old kite’; Simulation 2).

## Methods

### Model architecture and processing

We simulate N400 amplitudes on articles using the Sentence Gestalt (SG) model (Rabovsky et al., 2018; St. John & McClelland, 1990), as displayed in Figure 1.

### Environment and training

To train the SG model (Fig. 1), we used the same model environment and training parameters as for our previous simulations (described in detail in Rabovsky et al., 2018). The model environment consists of sentences (presented word by word at the input layer) such as ‘At breakfast, the girl eats an egg’ each paired with a corresponding event description, specifying who does what to whom in the described event, i.e., who is the agent of the event, what is the action in the event, what is the patient or object in the event, etc. Thus, the event specification consists of a set of pairs of thematic roles (e.g., agent, action, etc.) and their fillers (e.g., girl, eating, etc.). The pairs of sentences and corresponding event specifications are probabilistically generated online during training according to pre-specified constraints. After each presented word (represented by a word-specific unit at the input layer), the model is probed concerning all aspects of the event described by the sentence in the query network (see Fig. 1). Responding to a probe consists in completing a role-filler pair when probed with either a thematic role (i.e., agent, action, patient, location, or situation; each represented by an individual unit at the probe and output layer) or a filler of a thematic role (e.g., the girl, to eat, the eggs, etc.). For the filler concepts, we used feature-based semantic representations that were handcrafted to create graded similarities between concepts roughly corresponding to real world similarities (Rabovsky et al., 2018). For each response, the model’s activation at the output layer is compared with the correct output, the gradient of the cross-entropy error measure for each connection weight and bias term in the query network is back-propagated through this part of the network, and the corresponding weights and biases are adjusted accordingly. At the SG layer, the gradient of the cross-entropy error measure for each connection weight and bias term in the update network is collected for the responses on all the probes for each word before being back-propagated through this part of the network and adjusting the corresponding weights and biases.

For the current simulations, we intended to keep changes to the previous model environment and training to a minimum, while including the characteristics necessary to address N400 effects on articles. Specifically, for Simulation 1, the model environment was adjusted to include articles, and for Simulation 2, we further adjusted the environment to include adjectives in addition to the articles. As for our previous simulations, 10 independently initialized models (with initial weights randomly varying between +/− .05) were exposed to 800,000 probabilistically generated example sentences, and we used a learning rate of 0.00001 and momentum of .9 throughout.

#### Adjustments for Simulation 1

For Simulation 1, we added two input units, representing the two indefinite articles (‘a’ and ‘an’), which were not associated with specific semantic features at the output layer. We presented these indefinite articles during training at the sentence position prior to the objects in the sentences, such as e.g. ‘The man waters a pine’. Crucially, during training each of ten specific actions (e.g. ‘water’) is followed by either a high probability object (e.g., ‘pine’ with a probability of .7) or a low probability object (e.g., ‘oak’ with a probability of .3), and we constructed the training environment such that for each action, the high and low probability objects differ in terms of the appropriate article (e.g., ‘The man waters a pine’ vs. ‘… an oak’). Across all actions, both articles are linked equally often to the high versus low probability objects so that the articles do not differ in overall frequency. Importantly, after being presented with sentence beginnings such as ‘The man waters an…’ the trained model can predict that the sentence will not continue with the high probability continuation (‘pine’) but instead with the lower probability continuation (‘oak’; see Fig. 1b).

#### Adjustments for Simulation 2

For Simulation 2, we additionally added adjectives between the articles and the nouns. Specifically, two further input units were added, which represent two adjectives, each of which followed one of the indefinite articles (they can be thought of as ‘old’ and ‘new’ as in ‘an old x’/ ‘a new x’; please note that the units’ labels do not influence model behavior, but instead just serve to help the reader to map the roles of the units to the roles of words in natural sentences). As both indefinite articles occurred equally frequently across the training environment, both adjectives occurred equally frequently as well. Both of these adjectives were compatible with both the high cloze and the low cloze probability objects, so that addition of the adjectives removed the predictive value of the articles concerning the upcoming nouns.

Please note that these simulations aim to isolate and illustrate the influences of specific manipulations and specific aspects of the environment while minimizing changes to our previous training environment (Rabovsky et al., 2018), rather than capture the complexity and richness of natural language environments. While this controlled and synthetic approach to training has the advantage of being transparent concerning the factors influencing model behavior, it does not allow to simulate processing of specific stimulus sentences used in empirical experiments. Future work scaling up the model to large scale naturalistic training environments is required to achieve this goal, though it will come at the cost of reduced transparency.

### Simulations

Simulations 1 and 2 differed in terms of the training environment as described above, but did not differ in terms of the simulation experiments themselves, as the goal of including the second simulation is to demonstrate the influence of predictive relationships in the learning environment. For each of the simulations, we presented an agent (‘man’), followed by each of the ten specific actions (e.g., ‘water’) and, for the high cloze condition, the indefinite article compatible with the high probability object (e.g., ‘a’, which is compatible with the high probability continuation ‘pine’), and for the low cloze condition, the article compatible with the low probability object (e.g., ‘an’, which is compatible with the low probability continuation ‘oak’). It is important to note that for models exposed to the training environment for the first simulation, the articles were predictive of the upcoming nouns, while this predictive relationship was removed by the adjectives presented between articles and nouns in the training environment for the second simulation. For both simulations, there were 10 items per condition, and the model’s N400 correlate was computed as the summed magnitude of the difference in SG layer activation between presentation of the action (word *n*-1) and the article (word *n*), as illustrated in Figure 1b.

## Results

The model’s N400 correlate to the articles is displayed in Figures 2 and 3 (Supplementary Figures S1 and S2 show the results by item). Figure 2 displays the results of Simulation 1, where the articles provide reliable cues to meaning, while Figure 3 displays the results of Simulation 2, where the articles do not provide reliable cues. For both simulations, we used linear mixed model analyses (LMMs) implemented in the packages lme4 (Bates, Mächler, Bolker, & Walker, 2015) and lmerTest (Kuznetsova, Brockhoff, & Christensen, 2017) in R (www.r-project.org) to analyze influences of article cloze probability on the model’s N400 correlate. We entered a fixed effect for the article cloze probability factor (sum coding: −1 / +1) and added crossed random effects for subjects and items, with uncorrelated random intercepts and random slopes (for the cloze probability factor) varying across models and varying across items. (Adding random effects correlations yielded invalid estimates of −1, indicating overfitting.)

**Figure 2.**
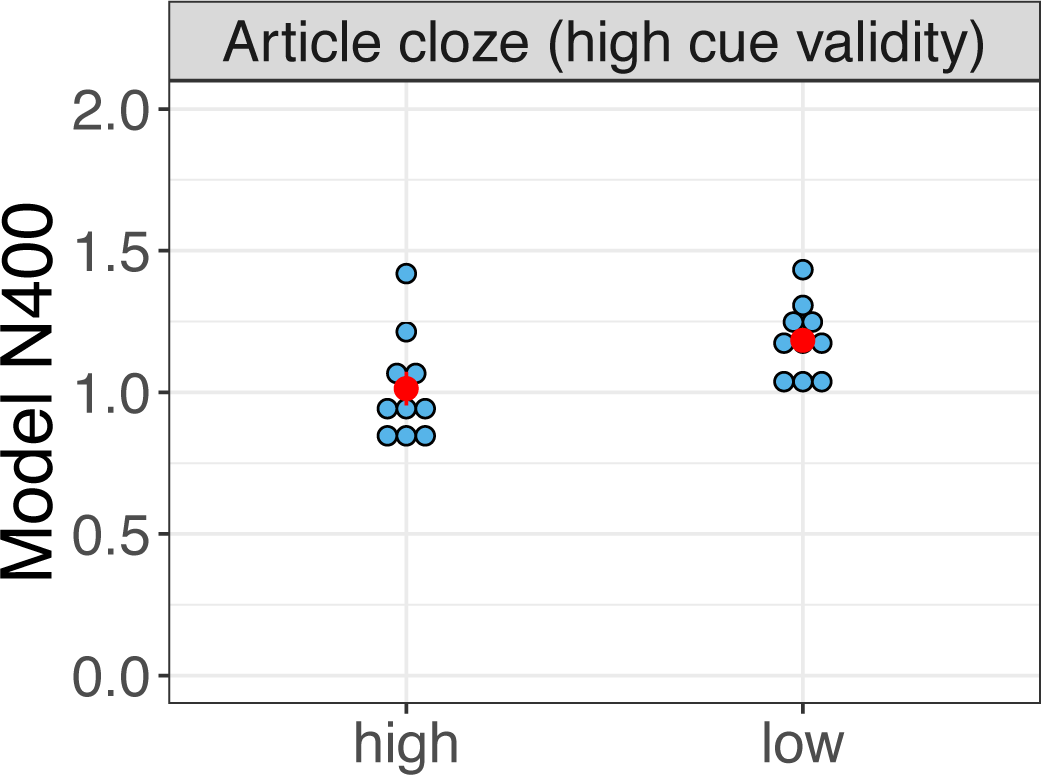
Displayed is the model’s N400 correlate as a function of article cloze probability after training on an environment where articles provide valid cues to meaning, because the articles are always directly followed by the nouns (in analogy to e.g., ‘a kite’/ ‘an airplane’; Simulation 1). Blue dots represent results for 10 independent runs of the model (averaged across 10 items per condition). Red dots represent condition means, +/− standard error of the mean (SEM) is represented by red error bars.

**Figure 3.**
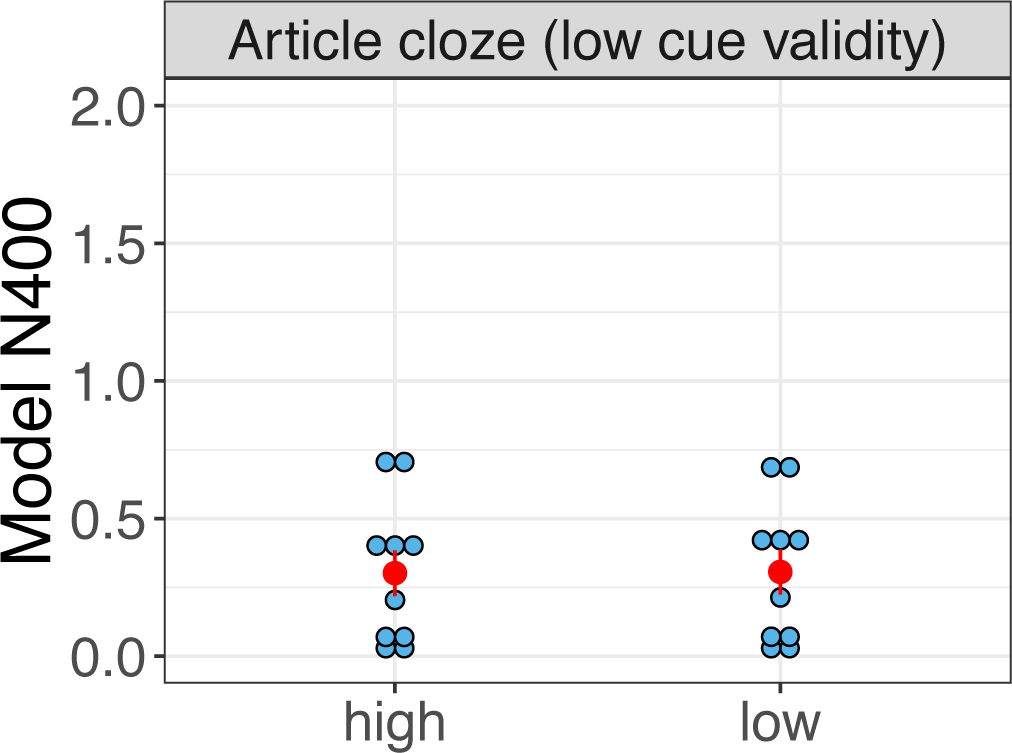
Displayed is the model’s N400 correlate as a function of article cloze probability after training on an environment where articles do not provide valid cues to meaning, because adjectives are presented between the articles and the nouns (in analogy to e.g., ‘an old kite’/ ‘a new airplane’; Simulation 2). Blue dots represent results for 10 independent runs of the model (averaged across 10 items per condition). Red dots represent condition means, +/− standard error of the mean (SEM) is represented by red error bars.

As can be seen In Figure 2, the model’s N400 correlate showed reliable article-induced modulations in Simulation 1, with larger semantic update (SU) for articles compatible with low cloze as compared to high cloze probability objects (*b* = 0.085, *SE* = 0.017, *t*_(180)_ = 4.998, *p* < 0.0001), in line with empirical studies (DeLong et al., 2005; Nicenboim et al., 2019). Please note that these effects are smaller than the simulated N400 effects for noun cloze probability, which we report in Rabovsky et al. (2018). It can also be seen that the effects on articles crucially depend on the predictive relationship between the articles and subsequent meaning, as they vanish in Simulation 2 when the cue validity of the articles is removed by the adjectives presented between articles and nouns during training (Fig. 3; *b* = 0.002, *SE* = 0.013, *t*_(180)_ = .174, *p* = 0.862).

## Discussion

The current study simulates article induced N400 effects (DeLong et al., 2005) as the change in a probabilistic and integrated representation of meaning, which corresponds to an implicit semantic prediction error, cued by encountering the articles. It additionally illustrates how these article induced N400 effects depend on the predictive relationship between articles and nouns in the linguistic environment.

Two issues seem to be worth discussing. First, the simulations provide a mechanistically explicit account of article induced N400 effects that does not depend on prediction at the level of word forms, but rather treats N400 amplitudes as the change in the conditional probabilities of semantic features cued by encountering the articles (Rabovsky et al., 2018; Rabovsky & McRae, 2014; Yan, Kuperberg, & Jaeger, 2017). This explanation differs from the most common interpretation of these effects (DeLong et al., 2005; Hagoort, 2017; Lau et al., 2008; Pickering & Garrod, 2013) and can, from the perspective implemented in the model, also be applied to related findings concerning article gender (see Kochari & Flecken, 2019 and Nicenboim et al., 2019 for recent reviews; see Fleur, Flecken, Rommers, & Nieuwland, 2019 for evidence consistent with the view that word form prediction may not be essential).

It is important to note that this does not mean that there is no probabilistic prediction of word forms during language comprehension. It simply suggests that if there is probabilistic prediction at the level of word forms, which may be expected if prediction is a fundamental aspect of brain function (Clark, 2013; Friston, 2005; McClelland, 1994; Rao & Ballard, 1999; Schultz, Dayan, & Montague, 1997), the consequences of probabilistic form prediction may be reflected in other (presumably earlier) ERP components (Gagl et al., 2018; Nieuwland, 2019).

The second issue is that one of the factors making it difficult to reliably observe article induced N400 effects in empirical studies is that the predictive relationship between indefinite articles and subsequent nouns is weakened by the fact that indefinite articles are not necessarily directly followed by nouns in natural language (Nieuwland et al., 2018). Here we demonstrate the impact of the articles’ cue validity by including simulations with very high versus very low cue validity, the assumption being that the cue validity in natural language presumably lies somewhere in between these extremes (according to the Corpus of Contemporary American English and British National Corpus, indefinite articles are directly followed by nouns in about a third of the cases; cited based on Nieuwland et al., 2018), thus contributing to the small size of the empirically observed effects (Nicenboim et al., 2019; Nieuwland et al., 2018).

A crucial point here seems to be that the same predictive comprehension system implemented in our model does produce article-induced N400 effects for situations with high but not low cue validity (corresponding to Simulations 1 and 2), and that this difference does not speak to the predictive nature of the system, but simply reflects the statistical regularities in the environment. The lack of an effect of article cloze probability in Simulation 2 is not due to the fact that the model does not predict, but rather due to the fact that the predictions of the model are not differentially adjusted based on the two articles (i.e., if the model predicts upon encountering the word ‘waters’ that the man will likely water a pine and less likely an oak, this prediction will not change upon encountering the indefinite article). Also note that because our model does not probabilistically predict the next word, but rather predicts integrated sentence meaning, the update to the model’s probabilistic representation does not necessarily depend on whether an expected noun is presented immediately or at a later point in the sentence, as long as new incoming evidence (e.g. an adjective presented in between) is still consistent with this expected noun.

In conclusion, the current simulations provide a mechanistically explicit account of article induced N400 effects as reflecting the change in a probabilistic representation of meaning corresponding to an implicit semantic prediction error, cued by the encountered articles depending on their cue validity. This account is in line with the view that the brain probabilistically predicts upcoming input based on the experience of statistical regularities in the environment (Clark, 2013; Friston, 2005; McClelland, 1994; Rao & Ballard, 1999; Schultz et al., 1997), and that the prediction error or Bayesian surprise at the level of meaning is reflected in N400 amplitudes (Kuperberg & Jaeger, 2016; Rabovsky et al., 2018; Rabovsky & McRae, 2014).

## Acknowledgments

This study was supported by an Einstein Prize Postdoctoral Stipend and a Visiting Fellowship at the Neurobiology of Language Department at the Max Planck Institute for Psycholinguistics (Nijmegen) to Milena Rabovsky.

## Supplementary Figures

**Supplementary Figure S1.**
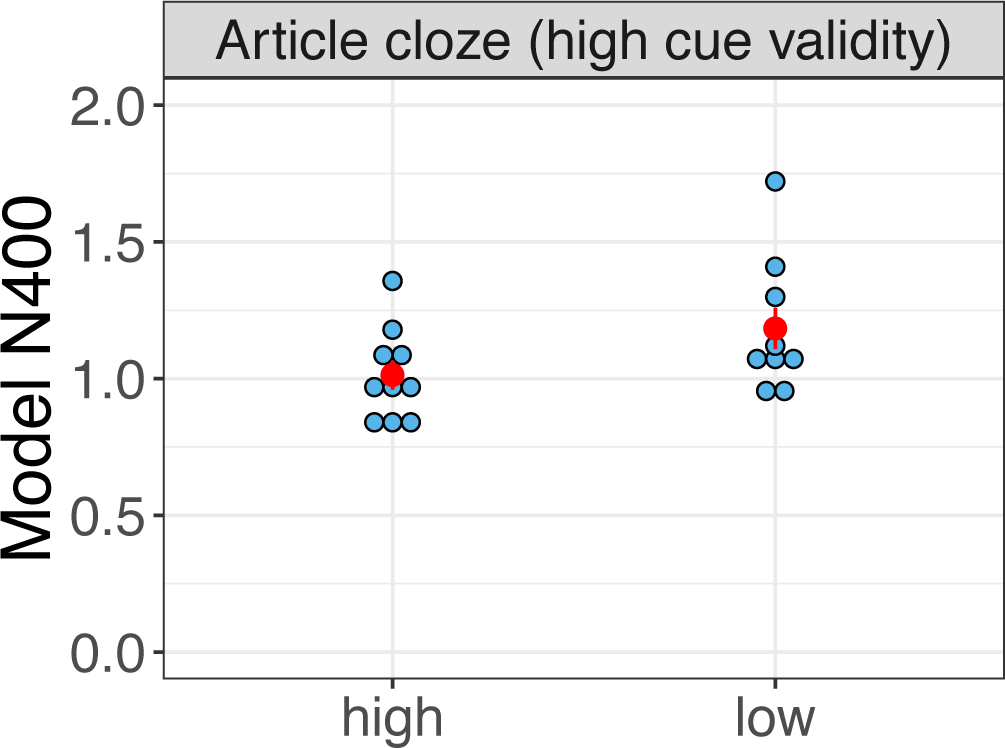
Displayed is the model’s N400 correlate (by item) as a function of article cloze probability after training on an environment where articles provide valid cues to meaning, because the articles are always directly followed by the nouns (in analogy to e.g., ‘a kite’/ ‘an airplane’; Simulation 1). Blue dots represent results for 10 items averaged across 10 independent runs of the model. Red dots represent condition means, +/− standard error of the mean (SEM) is represented by red error bars.

**Supplementary Figure S2.**
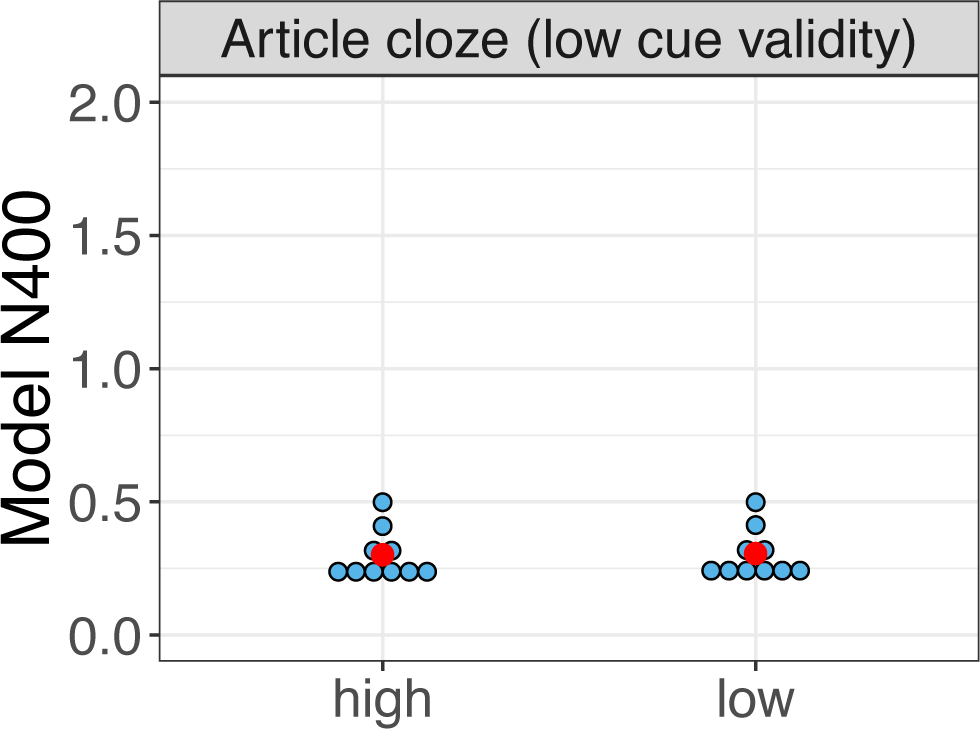
Displayed is the model’s N400 correlate (by item) as a function of article cloze probability after training on an environment where articles do not provide valid cues to meaning, because adjectives are presented between the articles and the nouns (in analogy to e.g., ‘an old kite’/ ‘a new airplane’; Simulation 2). Blue dots represent results for 10 items averaged across 10 independent runs of the model. Red dots represent condition means, +/− standard error of the mean (SEM) is represented by red error bars.

## References

Altmann, G. T. M., & Mirković, J. (2009). Incrementality and prediction in human sentence processing. Cognitive Science, 33(4), 583–609. http://doi.org/10.1111/j.1551-6709.2009.01022.x

Baggio, G., & Hagoort, P. (2011). The balance between memory and unification in semantics: A dynamic account of the N400. Language and Cognitive Processes, 26(9), 1338–1367. http://doi.org/10.1080/01690965.2010.542671

Bates, D., Mächler, M., Bolker, B., & Walker, S. (2015). Fitting Linear Mixed-Effects Models using lme4. Journal of Statistical Software, 67(1), 1–48. http://doi.org/10.18637/jss.v067.i01

Brown, C., & Hagoort, P. (1993). The processing nature of the N400: Evidence from masked priming. Journal of Cognitive Neuroscience, 5(1), 34–44. http://doi.org/10.1162/jocn.1993.5.1.34

Clark, A. (2013). Whatever next? Predictive brains, situated agents, and the future of cognitive science. Behavioral and Brain Sciences, 36(3), 181–204. http://doi.org/10.1017/S0140525X12000477

DeLong, K. A., Urbach, T. P., & Kutas, M. (2005). Probabilistic word pre-activation during language comprehension inferred from electrical brain activity. Nature Neuroscience, 8(8), 1117–1121. http://doi.org/10.1038/nn1504

DeLong, K. A., Urbach, T. P., & Kutas, M. (2017). Is there a replication crisis? Perhaps. Is this an example? No: a commentary on Ito, Martin, and Nieuwland (2016). Language, Cognition and Neuroscience, 32(8), 966–973. http://doi.org/10.1080/23273798.2017.1279339

Fleur, D. S., Flecken, M., Rommers, J., & Nieuwland, M. S. (2019). Definitely saw it coming? An ERP study on the role of article gender and definiteness in predictive processing Damien. BioRxiv. Retrieved from https://doi.org/10.1101/563783

Friston, K. (2005). A theory of cortical responses. Philosophical Transactions of the Royal Society of London. Series B, Biological Sciences, 360(1456), 815–36. http://doi.org/10.1098/rstb.2005.1622

Gagl, B., Sassenhagen, J., Haan, S., Gregorova, K., Richlan, F., & Fiebach, C. J. (2018). Visual word recognition relies on an orthographic prediction error signal. BioRxiv. Retrieved from http://dx.doi.org/10.1101/431726

Hagoort, P. (2017). The core and beyond in the language-ready brain. Neuroscience and Biobehavioral Reviews, 81, 194–204. http://doi.org/10.1016/j.neubiorev.2017.01.048

Ito, A., Martin, A. E., & Nieuwland, M. S. (2017a). How robust are prediction effects in languagecomprehension? Failure to replicate article-elicitedN400 effects. Language, Cognition, and Neuroscience, 32(8), 954–965. http://doi.org/10.1016/j.jml.2013.08.001

Ito, A., Martin, A. E., & Nieuwland, M. S. (2017b). Why the A/AN prediction effect may be hard to replicate: a rebuttal to Delong, Urbach, and Kutas (2017). Language, Cognition and Neuroscience, 32(8), 974–983. http://doi.org/10.1080/23273798.2017.1323112

Kochari, A. R., & Flecken, M. (2019). Lexical prediction in language comprehension: a replication study of grammatical gender effects in Dutch. Language, Cognition and Neuroscience, 34(2), 239–253. http://doi.org/10.1080/23273798.2018.1524500

Kuperberg, G. R., & Jaeger, T. F. (2016). What do we mean by prediction in language comprehension? Language, Cognition and Neuroscience, 31(1), 32–59. http://doi.org/10.1080/23273798.2015.1102299

Kutas, M., & Federmeier, K. D. (2000). Electrophysiology reveals semantic memory use in language comprehension. Trends in Cognitive Sciences. http://doi.org/10.1016/S1364-6613(00)01560-6

Kutas, M., & Federmeier, K. D. (2011). Thirty years and counting: finding meaning in the N400 component of the event-related brain potential (ERP). Annual Review of Psychology, 62(August), 621–647. http://doi.org/10.1146/annurev.psych.093008.131123

Kutas, M., & Hillyard, S. A. (1984). Brain potentials during reading reflect word expectancy and semantic association. Nature, 307, 101–103.

Kuznetsova, A., Brockhoff, P. B., & Christensen, R. H. B. (2017). lmerTest Package: Tests in Linear Mixed Effects Models. Journal of Statistical Software, 82(13), 1–26. http://doi.org/10.18637/jss.v082.i13

Lau, E. F., Phillips, C., & Poeppel, D. (2008). A cortical network for semantics: (de)constructing the N400. Nature Reviews Neuroscience, 9(12), 920–933. http://doi.org/Doi10.1038/Nrn2532

McClelland, J. L. (1994). The interaction of nature and nurture in development: A parallel distributed processing perspective. International Perspectives on Psychological Science, Volume 1: Leading Themes.

Nicenboim, B., Vasishth, S., & Rösler, F. (2019). Are words pre-activated probabilistically during sentence comprehension? Evidence from new data and a Bayesian random-effects meta-analysis using publicly available data. *PsyArXiv*. Retrieved from https://doi.org/10.31234/osf.io/2atrh%0A

Nieuwland, M. S. (2019). Do ‘early’ brain responses reveal word form prediction during language comprehension? A critical review. Neuroscience and Biobehavioral Reviews, 96(May 2018), 367–400. http://doi.org/10.1016/j.neubiorev.2018.11.019

Nieuwland, M. S., Politze-Ahles, S., Heyselaar, E., Segaer, K., Bartolozzi, F., Kogan, V.,… Huettig, F. (2018). Large-scale replication study reveals a limit on probabilistic prediction in language. ELIFE, (5030732), 1–24. http://doi.org/10.7554/eLife.33468

Pickering, M. J., & Garrod, S. (2013). An integrated theory of language production and comprehension. Behavioral and Brain Sciences, 36(04), 329–347. http://doi.org/10.1017/s0140525x12001495

Rabovsky, M., Hansen, S. S., & McClelland, J. L. (2018). Modelling the N400 brain potential as change in a probabilistic representation of meaning. Nature Human Behaviour, 2, 693–705. http://doi.org/10.1038/s41562-018-0406-4

Rabovsky, M., & McRae, K. (2014). Simulating the N400 ERP component as semantic network error: Insights from a feature-based connectionist attractor model of word meaning. Cognition, 132(1), 68–89. http://doi.org/10.1016/j.cognition.2014.03.010

Rao, R. P. N., & Ballard, D. H. (1999). Predictive coding in the visual cortex: a functional interpretation of some extra-classical receptive-field effects. Nature, 2(1), 79–87.

Schultz, W., Dayan, P., & Montague, P. R. (1997). A neural substrate of prediction and reward. Science, 275(5306), 1593–1599. http://doi.org/10.1126/science.275.5306.1593

St. John, M. F., & McClelland, J. L. (1990). Learning and applying contextual constraints in sentence comprehension. Artificial Intelligence, 46(1–2), 217–257. http://doi.org/10.1016/0004-3702(90)90008-N

Yan, S., Kuperberg, G. R., & Jaeger, T. F. (2017). Prediction (or not) during language processing. A commentary on Nieuwland et al. (2017) and DeLong et al. (2005) Shaorong. BioRxiv. Retrieved from http://dx.doi.org/10.1101/143750

